# Anti-BP180 Autoantibodies Are Present in Stroke and Recognize Human Cutaneous BP180 and BP180-NC16A

**DOI:** 10.1101/313692

**Authors:** Yanan Wang, Xuming Mao, Di Wang, Christoph M. Hammers, Aimee S. Payne, Yiman Wang, Hongzhong Jin, Bin Peng, Li Li

**Author notes:** **Corresponding author:** Li Li, Department of Dermatology, Peking Union Medical College Hospital, Chinese Academy Medical Science, No. 1 Shuai Fu Yuan Street, Beijing, China, 100730. Bin Peng, Department of Neurology, Peking Union Medical College Hospital, Chinese Academy Medical Science, No. 1 Shuai Fu Yuan Street, Beijing, China, 100730. **Funding sources**: This study was supported by National Natural Science Foundation of China (81371731), Milstein Medical Asian American Partnership foundation (2017, Dermatology) and the Education Reform Projects of Peking Union Medical College (No. 2016zlgc0106). IRB approval status: Reviewed and approved by IRB members of Peking Union Medical College Hospital, CAMS & PUMC, Protocol Number: S-300. **Conflicts of Interest**: None declared.

## Abstract

**Background:** Current evidence has revealed a significant association between bullous pemphigoid (BP) and neurological diseases (ND), including stroke, but the incidence of BP autoantibodies in patients with stroke has not previously been investigated.

**Objective:** Our study aims to assess BP antigen-specific antibodies in stroke patients.

**Methods:** 100 patients with stroke and 100 healthy controls were randomly selected to measure anti-BP180/230 IgG autoantibodies by enzyme-linked immunosorbent assay (ELISA), salt split indirect immunofluorescence (IIF) and immunoblotting against human cutaneous BP180 and BP180-NC16A.

**Results:** Anti-BP180 autoantibodies were found in 14(14.0%) patients with stroke and 5(5.0 %) of controls by ELISA (p<0.05). Sera from 13(13.0%) patients with stroke and 3(3.0 %) controls reacted with 180-kDa proteins from human cutis extract (p<0.05). 11(11.0%) of stroke and 2(2.0 %) of control sera recognized the human recombinant full length BP180 and NC16A (p<0.05). The anti-BP180-positive patients were significantly younger than the negative patients in stroke (p<0.001).

**Limitations:** Longitudinal changes in antibody titers and long-term clinical outcome for a long duration were not fully investigated.

**Conclusion:** Development of anti-BP180 autoantibodies occur at a higher frequency after stroke, suggesting BP180 as a shared autoantigen in stroke with BP and providing novel insights into BP pathogenesis in aging.

## INTRODUCTION

Bullous pemphigoid (BP) is a common autoimmune blistering skin disorder most commonly found in the elderly^1^. BP is characterized by circulating and tissue-bound autoantibodies directed against two hemidesmosomal components: the transmembrane BP180 (collagen XVII, BPAG2) protein, and the plakin family protein BP230 (BPAG1)^2^, which are localized at the basement membrane zone (BMZ) of skin, as well as central nervous system (CNS) and other tissues. In human brain, BP180 is expressed in the hypoglossal nucleus, oculomotor nucleus, and pyramidal cells of the hippocampal regions^3^. Anti-BP180 immunoglobulin G (IgG) autoantibodies contribute to blister formation and correlate with disease activity, particularly at the time of diagnosis and at disease flare^4^. The immunodominant epitopes of BP180 are localized in the extracellular non-collagenous 16A (NC16A) domain^5–7^. In contrast, anti-BP230 IgG autoantibodies, which are also found in the majority of BP patients, are considered of minor pathogenic relevance^8^, most likely due to its intracellular localization.

The significant association between BP and neurological diseases (ND) has been fully supported by a series of previous studies^9–11^. Patients older than 80 years with ND were 10 times more likely to have BP than those without^11^, and the most frequently associated conditions were cerebral stroke in men and dementia in women. Stroke, a life-threatening condition and the second cause of death worldwide, is one of the most common forms of ND. In a study including 12,607 patients with first-ever stroke, 38 (0.3%) patients developed BP in a median of 3.5 years, while only 8 people (0.06%) had BP in a median of 3.7 years in the control group^12^. Additionally, there was a 2-fold increase in risk of developing BP in those with acute ischemic stroke in a population-based case-control study in the UK^9^. In the current study, we aimed to determine if there is a higher incidence of BP specific autoantibodies in stroke patients, which may underlie the increased incidence of BP in this population.

## RESULTS

### BP180 and BP230 autoantibodies and short-term clinical outcomes in patients with stroke

Sera from patients with stroke (n=100) (including cerebral infarction and cerebral hemorrhage) and healthy controls (n=100) were collected to examine anti-BP180/230 IgG antibodies by ELISA (cut-off value >9 U/ml). The positive rate of anti-BP180 antibody in the stroke cohort (14, 14.0%) was significantly higher than that in controls (5, 5.0%) (P=0.03) (Table 1, 3, 4). All anti-BP180 IgG positive patients (14 stroke samples and 5 healthy controls) were further examined by immunoblotting against human cutis extract, human recombinant full length BP180 and human recombinant NC16A (Table 1, 3, 4). Sera from 13(13.0%) stroke patients and 3 (3.0 %) healthy controls reacted with 180-kDa proteins from the human cutis extract (P=0.016) (Fig 1a). Sera from 11 (11.0%) patients with stroke and 2(2.0 %) healthy controls recognized both of the human recombinant full length BP180 (P=0.018) (Fig 1b) and human recombinant NC16A (P=0.018) (Fig 1c). Anti-BP180 positive sera were further tested by salt-split IIF, and only one patient with stroke revealed that IgG antibody binding on the epidermal side of BMZ (Fig 2).

**Table 1.** Antibody positive rates between stroke and control

|  | Stroke<br>(n = 100) | Control<br>(n = 100) | P value |
| --- | --- | --- | --- |
| ELISA | 14(14.0%) | 5(5.0%) | 0.03* |
| Immunoblotting |  |  |  |
| Human cutis extract | 13(13.0%) | 3(3.0%) | 0.016* |
| Human recombinant full length BP180 | 11(11.0%) | 2(2.0%) | 0.018* |
| Human recombinant NC16A | 11(11.0%) | 2(2.0%) | 0.018* |
| Salt split IIF | 1(1.0%) | 0(0.0%) | 0.316 |

ELISA: enzyme-linked immunosorbent assay, IIF: indirect immunofluorescence.

**Figure 1.**
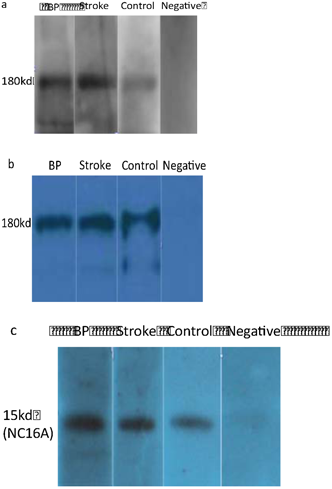
(a) A patient with bullous pemphigoid (BP), a stroke patient and a control patient recognized a 180-kDa protein from human cutis extract. (b) A patient with BP, a stroke patient and a control patient recognized human recombinant full length BP180. (c) A patient with BP, a stroke patient and a control patient recognized human recombinant BP180-NC16A.

**Figure 2.**
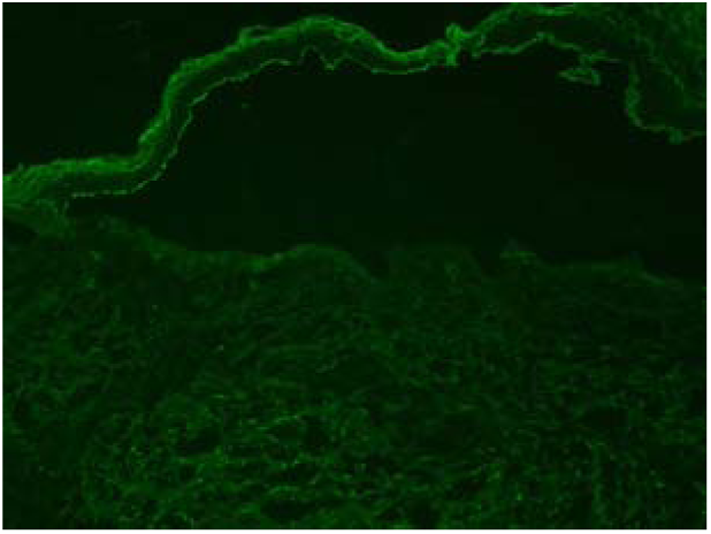
A patient with stroke showed IgG antibody deposition on the epidermal side of BMZ.

The positive rate of anti-BP230 antibodies has no statistical difference between the stroke (14, 14.0%) and control groups (15, 15%) in the ELISA assay. Statistical analysis showed there was no significant difference in sex and age between the stroke and control group (Table 2). For complications, significant differences were found in cardiovascular diseases (P<0.01) and tumor (P<0.01) (Table 2) between the two groups. Using logistic regression analysis and the Hosmer-Lemeshow test, cardiovascular disease and stroke had a positive correlation (p<0.01), while tumor and stroke had a negative correlation (P<0.01). In contrast, according to bivariate Kendall coefficient analysis, anti-BP180 antibody positivity was found to be an independent factor that had no significant correlation with cardiovascular disease or tumor.

**Table 2.** Analysis of age, sex and complications between stroke and control

|  | Stroke<br>(n = 100) | Control<br>(n = 100) | OR (95% CI) | P value |
| --- | --- | --- | --- | --- |
| Male, n (%) | 69 (69.0) | 74 (74.0) |  | 0.434 |
| Age, n (%) |  |  |  |  |
| < 60 yr | 30 (30.0) | 30 (30.0) |  | >0.999 |
| 60 – 69 yr | 30 (30.0) | 32 (32.0) |  | 0.760 |
| 70 – 79 yr | 25 (25.0) | 20 (20.0) |  | 0.397 |
| ≥ 80 yr | 15 (15.0) | 19 (19.0) |  | 0.451 |
| Complications, n (%) |  |  |  |  |
| None | 13 (13.0) | 20 (20.0) | 1.673 (0.781-3.583) | 0.182 |
| Cardiovascular disease | 77 (77.0) | 46 (46.0) | 0.254 (0.138-0.468) | <0.001* |
| Hyperlipidemia | 29 (29.0) | 19 (19.0) | 0.574 (0.297-1.112) | 0.098 |
| Diabetes | 21 (21.0) | 30 (30.0) | 1.612 (0.847-3.069) | 0.144 |
| Chronic kidney disease | 7 (7.0) | 2 (2.0) | 0.271 (0.055-1.339) | 0.170 |
| Other ND | 3 (3.0) | 3 (3.0) | 1.00 (0.197-5.078) | >0.999 |
| Digestive system disease | 2 (2.0) | 5 (5.0) | 2.579 (0.488-13.617) | 0.445 |
| Skin disease | 0 (0) | 4 (4.0) | 1.042 (1.001-1.084) | 0.121 |
| Tumor | 3 (3.0) | 18 (18.0) | 7.098(2.019-24.950) | <0.001* |
| BP180 (+), n (%) | 14 (14.0) | 5 (5.0) | 0.323 (0.112-0.935) | 0.030* |

\* Significant differences

### Analysis of anti-BP180 positive patients in stroke group and control group

Anti-BP180 autoantibody titers were significantly higher in the stroke group (19.2±6.07 U/ml) compared to those of control group (12.2±2.39 U/ml; P=0.024) (Table 3, 4). The medical records of anti-BP180 positive patients and controls were reviewed and all patients were followed up until October 2017. In the 1-3-year follow-up period, neither stroke patients nor the controls developed BP-like skin lesions. The 1-3-year survival rate of anti BP180 positive patients with stroke and control were both 100%.

**Table 3.** Characteristics and Immunological results of anti-BP180 positive patients of stroke

| Age | Sex | Diagnosis | BP180 U/ml | Immunoblotting |  |  | Salt-split IIF |
| --- | --- | --- | --- | --- | --- | --- | --- |
|  |  |  |  | Human cutis extract | BP180 | NC16A |  |
| 79 | M | CI and CH | 17 | + | + | + | - |
| 84 | M | CI | 19 | + | - | - | - |
| 54 | M | CI | 23 | + | + | + | - |
| 45 | M | CI | 19 | + | - | - | - |
| 56 | M | CI | 15 | + | + | + | - |
| 69 | M | CI | 30 | + | + | + | - |
| 62 | F | CI | 17 | + | + | + | IgG |
| 53 | M | CH | 23 | + | + | + | - |
| 57 | M | CI | 22 | + | + | + | - |
| 61 | F | CI | 10 | + | + | + | - |
| 64 | M | CI | 18 | + | + | + | - |
| 55 | M | CI | 30 | + | + | + | - |
| 52 | F | CI | 10 | - | - | - | - |
| 51 | M | CI | 16 | + | + | + | - |

CI: Cerebral infarction, CH: Cerebral hemorrhage, IIF: Indirect immunofluorescence, +: Positive, -: Negative. BP180: Human recombinant full length cutaneous BP180, NC16A: Human recombinant cutaneous BP180-NC16A

**Table 4.** Characteristics and Immunological results of anti-BP180 positive patients of control

| Age | Sex | BP180 U/ml | Immunoblotting |  |  | Salt split IIF |
| --- | --- | --- | --- | --- | --- | --- |
|  |  |  | Human cutis extract | BP180 | NC16A |  |
| 65 | M | 11 | + | + | - | - |
| 71 | M | 11 | + | + | - | - |
| 71 | M | 10 | - | - | + | - |
| 70 | M | 16 | - | - | - | - |
| 61 | F | 13 | + | - | + | - |

BP180: Human recombinant full length cutaneous BP180, NC16A: Human recombinant cutaneous BP180-NC16A, IIF: Indirect immunofluorescence, +: Positive, -: Negative.

### Comparison between anti-BP180 positive stroke patients and anti-BP180 negative stroke patients

According to statistical analysis, the average age of the anti-BP180 positive group (60.1 yrs) was significantly lower than that of the anti-BP180 negative group (69.0 yrs; P<0.001). Among them, the proportion of patients younger than 60 years in the anti-BP180 positive group (8/14, 57.1%) was significantly higher than that of the anti-BP180 negative patients (19/86, 24.4%; P=0.006). The duration of follow up after first stroke attack of BP180 positive group (7.0±2.94 yrs) was significantly shorter than that of the anti-BP180 negative group (10.4±6.05 yrs; P<0.001). There was no significant difference in sex, complications, and stroke attack times between the two groups (Table 5).

**Table 5.** Demographic characteristics of anti-BP180 positive/negative patients of stroke

| Variable | BP180 (+)<br>(n = 14) | BP180 (-)<br>(n = 86) | P value |
| --- | --- | --- | --- |
| Male, n (%) | 11 (78.6) | 58(67.4) | 0.540 |
| Age, mean±sd | 60.1±11.20 | 69.0±11.67 | <0.001* |
| < 60 yr, n (%) | 8 (57.1) | 19 (24.4) | 0.006* |
| 60 – 70 yr, n (%) | 4 (28.6) | 36 (46.2) | 0.395 |
| ≥ 75 yr, n (%) | 2 (14.3) | 23 (29.5) | 0.508 |
| Stroke attack times ≥ 2 , n (%) | 6 (42.9) | 21 (24.4) | 0.194 |
| Duration after first attack ≥ 1yr, n (%) | 9 (64.3) | 49 (57.0) | 0.607 |
| Duration after first attack (y), median | 7.0±2.94 | 10.4±6.05 | <0.001* |
| Complications, n (%) |  |  |  |
| No Complications | 4 (28.6) | 9 (10.5) | 0.077 |
| Cardiovascular disease | 10 (71.4) | 67 (77.9) | 0.704 |
| Hyperlipidemia | 4 (28.6) | 25 (29.1) | > 0.999 |
| Diabetes | 2 (14.2) | 19 (22.1) | 0.728 |
| Chronic kidney disease | 1 (7.1) | 6 (7.0) | > 0.999 |
| Other nervous system diseases | 0 (0) | 3 (3.5) | > 0.999 |
| Digestive system disease | 0 (0) | 2 (2.3) | > 0.999 |
| Tumor | 0 (0) | 3 (3.5) | > 0.999 |

BP180: Human recombinant full length cutaneous BP180

## DISCUSSION

Previously we have shown that a significant proportion of serum samples obtained from BP+ND patients could react with BP antigens of the human brain and skin, confirming the existence of BP antigens in the brain of ND patients^13,14^. In this work, we measured BP autoantibodies in 100 Chinese patients with stroke, demonstrating that anti-BP180 autoantibodies are at a higher frequency in stroke patients and that they recognize human cutaneous BP180 and BP180-NC16A. The positive rate of BP180 antibody in 337 healthy Americans was 3.7% (11/297) by ELISA^15^, which was similar to our control group (5/100, 5%). These observations raise the possibility that BP 180 acts as a shared autoantigen in both stroke and BP. We speculate that damage or alterations in the human CNS during the course of stroke could expose the neuronal isoform of BP180, thus triggering an immune reaction that, along with the immunological cross-reactions, may eventually result in cutaneous damage^13,14^.

In our study, only one stroke patient positive both for BP180 ELISA and immunoblotting (human BP180 and BP180-NC16A) displayed binding of IgG autoantibodies to the epidermal side of the BMZ, consistent with the work by Messingham et al^16^ and Kokkonen et al^17^ in Alzheimer’s disease and Parkinson’s disease. This might be at least partly explained by some additional triggers required for BP antibody development^17^.

Notably, we found that anti-BP180 positive stroke patients (60.1 yrs) were significantly younger than anti-BP180 negative stroke patients (69.0 yrs; P<0.001), suggesting young age might be a risk factor for stroke patients to develop BP. The proportion of patients younger than 60 years in anti-BP180 positive patients (8/14, 57.1%) was significantly higher than that of anti-BP180 negative patients (19/86, 24.4%; P=0.006) (Table 5). The duration of follow up after first stroke attack of anti-BP180 positive patients (7.0±2.94y) was significantly shorter than that of anti-BP180 negative patients (10.4±6.05y; P<0.001), further indicating that younger stroke patients with shorter duration after first attack are more likely to develop BP antibodies. This may be related to the immune responses in the early stage of stroke and down regulation of immune responses in the late stage of stroke^19^.

During our follow-ups, neither the anti-BP180/230 positive stroke patients nor the controls exhibited BP-like skin lesions. In accordance with this result, another group showed none of the anti-BP180/230 antibody positive individuals in their study revealed BP-like skin lesions^15^. Moreover, Kokkonen et al showed BP180 autoantibodies were found in 18% of patients with Alzheimer’s disease and 3% of controls (P = 0.019), while none of them had BP-like lesions^17^. The reason why anti-BP antibody positive patients had no BP-like lesions may be due to a few possibilities: First, titers of anti-BPAG antibodies in these subjects may be too low to cause cutaneous lesions. Second, some patients may be misdiagnosed because of atypical lesions. In about 20% of BP patients, atypical lesions arise as prurigo-like nodules, intertrigo-like pemphigoid, and localized forms. Finally, in about 10–20% of BP patients, disease onset is preceded by a prodromal phase of weeks to months with pruritus, excoriations, and eczematous lesions, and some patients never develop blisters^20^. Future studies with longer clinical follow-ups will be necessary to clarify the association of BP autoantibody development with overt clinical disease.

The positive rate of anti-BP180 ELISA in stroke and control groups was a little higher than that of immunoblotting in our study (Table 1). A meta-analysis described ELISA as a quantitative test with high sensitivity and specificity (87% and 98%–100%, respectively) for diagnosis of BP^21^, while immunoblotting was a semi-quantitative test. Therefore the ELISA positive stroke patient with low titers may be negative for immunoblotting. It is also important to note that for BP180, 7.8 % of BP sera react exclusively to regions of BP180 outside of NC16A region, which could not be identified by using the commercially available BP180 NC16A ELISA test. Thus, a negative test should be closely followed up with DIF and/or IIF^22^. There were no ELISA negative patients that were positive for IIF in our study.

Anti-BP180 autoantibody values were significantly higher in the stroke cohort (19.2±6.07 U/ml) compared to those in the control group by ELISA (12.2±2.39 U/ml; P=0.024) (Table 3,4). As the anti-BP180 antibodies are correlated with the activity of BP, we consider that the difference between the two groups is of clinical significance and may herald the development of BP in the future. Jedlickova et al found that male stroke patients were more likely to have BP^11^. In our study the proportion of male anti-BP180 positive patients (11/14, 84.6%) was slightly higher than that of controls (4/5, 80.0%; P>0.999), albeit with no significant difference. A larger sample size might be required for a statistical significance.

We conclude that anti-BP180 autoantibody occur at a higher frequency in stroke patients than age-and sex-matched controls suggesting that BP 180 could serve as a shared autoantigen in both stroke and BP and may underlie the epidemiologic association of these two conditions. Our study explained partially the mechanism of BP associated with ND and provided a theoretical basis for the prevention of BP in patients with stroke. Future studies using longer-term clinical follow up as well as animal or cell models may help to further clarify why BP is significantly associated with aging and neurologic diseases.

## MATERIALS AND METHODS

### Patient samples

This study was approved by the Ethical Committee of Peking Union Medical College Hospital and informed consents were obtained from all patients and normal individuals. Patients with stroke (cerebral infarction and cerebral hemorrhage) were from the department of Neurology of Peking Union Medical College Hospital during July 2014 to October 2016. All the patients with stroke were diagnosed in the department of Neurology. The age and sex matched control group was from patients attending the hospital for surgery from 2014 to 2016, and the patients with neurological diseases (other cerebrovascular disease, Parkinson’s disease, dementia, multiple sclerosis and amyotrophic lateral sclerosis) and skin diseases (bullous skin disease, dermatitis and eczema) were excluded after reviewing medical records. There was no significant difference in sex and age between the stroke and the control cohorts. The patients with BP, visiting Peking Union Medical College Hospital between 2013 and 2016, were diagnosed based on typical clinical, histopathological features and IgG and/or C3 deposition to BMZ in indirect immunofluorescence. Normal human foreskin tissues were taken from patients receiving circumcision in the Department of Urology.

BP180 and BP230 enzyme-linked immunosorbent analysis (ELISA)

Anti-BP180/230 IgG autoantibodies in the sera samples of patients and healthy controls were detected by commercially available ELISAs of human BP180 (NC16A domains) IgG (MEASACUP BP180, MBL, Japan) and BP230 N-and C-terminal domains IgG (BP230 ELISA kit, MBL, Japan) according to the manufacturer’s instruction (based on a cut-off value >9 U/ml).

### Immunoblotting

Protein extract preparations, polyacrylamide gel electrophoresis and immunoblotting were performed as previously described^13^. Briefly, human skin samples were subjected to Cell Lysis Solution (Thermo Fisher Scientific, Massachusetts, U.S.A). Following homogenization, ice-incubation (30 minutes) and centrifugation (12000rpm, 4°C, 15 minutes), the supernatant was collected and added with loading buffer. Human full length BP180/NC16A gene fragments were synthesized and subcloned to the pcDNA3.1 mammalian expression vector. HEK 293 cells were transiently transfected with the plasmids and lipofectamine (Life Technologies, Carlsbad, CA, USA) per the manufacturer’s instructions, followed by lysis of the cells in a lysis buffer (50mM Tris (PH8.0), 300mM NaCl, 1% Triton X-100, 1mM DTT, 5% glycerol). The proteins in lysates were purified. Expression of the target proteins were confirmed by western-blot (anti-Flag tag antibody) and quantified by the protein quantification assay kit with final concentrations of 600-1000μg/ml. Proteins were subjected to 8% SDS-PAGE gels under denaturing conditions, transferred onto a PVDF membrane (Thermo Fisher Scientific, Massachusetts, U.S.A), and incubated with human serum samples as the primary antibody and anti-human lgG-HRP (Abcam, Cambridge, Britain) as the secondary antibody. Membranes were developed with detection solution (Merck KGaA, Darmstadt, Germany) and the protein side of the membrane was exposed to image analysis system (Tanon, Shanghai, China). An anti-human Collagen XVII antibody (Abcam, Cambridge, Britain) (primary antibody) and goat anti-rabbit IgG H&L (HRP) antibodies (Abcam, Cambridge, Britain) (secondary antibody) were used as the positive control to detect BP180-protein in immunoblots.

### Salt split indirect immunofluorescence (IIF)

5-μm frozen non-fixed sections of human skin (treated with 1 M NaCl) were blocked (1% BSA in PBS) and human sera in 1:4 to 1:320 dilutions were used as primary antibodies. Rabbit anti-human IgG-FITC (Abcam, Cambridge, Britain) was used as the secondary antibody. The skin sections were then mounted with glycerol/PBS (2:1, pH 9.0) and observed under a fluorescence microscope. A clear linear immunostaining on the BMZ was considered positive, while no fluorescence was considered negative.

### Statistical analysis

The data involved in the statistical analysis included qualitative analysis, such as BP180 antibody (positive/negative), sex (male/female), complication (yes/no) and frequency of attack (single/ multiple); and quantitative analysis (such as age, BP180/230 antibody titer). Specifically, qualitative data were statistically analyzed by chi square test and logistic regression analysis, and the quantitative data were statistically analyzed by t test and rank sum test.

## ACKNOWLEDGMENTS

We gratefully acknowledge the technical assistance of Nan Yang, PhD.

## Abbreviations

BP: Bullous pemphigoid
IIF: Indirect immunofluorescence
DIF: Direct immunofluorescence
ND: Neurological diseases
NC16A: Non-collagenous 16A
ELISA: Enzyme-linked immunosorbent assay
BMZ: Basement membrane zone
BBB: Blood–brain barrier
CNS: Central nervous system

